# Cullin3 promotes stem cell progeny differentiation by facilitating aPKC-directed asymmetric Numb localization

**DOI:** 10.1101/2020.07.29.227603

**Authors:** Hideyuki Komori, Noemi Rives-Quinto, Xu Han, Lucas Anhezini, Ari J. Esrig, James B. Skeath, Cheng-Yu Lee

## Abstract

Asymmetric segregation of Numb is a conserved mechanism for regulating Notch-mediated binary cell fate decisions; however, the mechanisms controlling Numb segregation remain unclear. Previous studies have proposed an “exclusion” model, suggesting that atypical protein kinase C (aPKC) negatively regulates Numb cortical localization. Here, we report that aPKC kinase activity positively promotes basal cortical Numb localization during asymmetric division of *Drosophila* neural stem cells (neuroblasts) and that Cullin 3 (Cul3) is required for aPKC-directed basal Numb localization. In *numb-* or *cul3*-mutant brains, decreased levels of Numb segregated into neuroblast progeny failed to downregulate Notch, leading to supernumerary neuroblast formation. Increased aPKC kinase activity suppressed supernumerary neuroblast formation by concentrating residual Numb protein at the basal cortex and in neuroblast progeny, whereas decreased aPKC function enhanced the supernumerary neuroblast phenotype by reducing basal Numb levels. We propose that aPKC and Cul3 promote basal Numb localization, which is required to downregulate Notch signaling and promote differentiation in neuroblast progeny.

## Introduction

Asymmetric cell division allows for the generation of two sibling cells with distinct identities and plays a key role during tissue development and organ homeostasis. Notch signaling provides a highly efficient mechanism for regulating binary cell fate decisions, as multiple mechanisms simultaneously regulate different layers of this pathway (Bray, 2016; Pinto-Teixeira and Desplan, 2014). One of these mechanisms is asymmetric segregation of the Notch antagonist Numb. Exclusive segregation of Numb into one of two progeny siblings following asymmetric cell division allows for the specification of one Notch^ON^ cell and one Notch^OFF^ cell (Knoblich et al., 1995; Zhong et al., 1996). During *Drosophila* larval brain neurogenesis, asymmetric segregation of Numb leads to downregulation of Notch signaling in neuroblast progeny, enabling the initiation of differentiation. The resultant progeny type depends on the neuroblast identity: type II neuroblasts generate intermediate neural progenitors (INPs), which have a limited proliferation potential, whereas type I neuroblasts generate ganglion mother cells (GMCs), which differentiate into a pair of neurons (Farnsworth and Doe, 2017; Homem et al., 2015; Janssens and Lee, 2014; Xiao et al., 2012). Despite these differences, both types of neuroblasts segregate Numb into their progeny, which antagonizes Notch to promote differentiation and prevent supernumerary neuroblast formation. Although critical for brain development, the mechanisms governing the precise localization of Numb during asymmetric cell division are not well understood.

Numb and the TRIM-NHL protein Brain tumor (Brat) asymmetrically localize to the basal cortex of mitotic neuroblasts and segregate into GMCs and immature INPs (Bello et al., 2006; Betschinger et al., 2006; Lee et al., 2006a; Lee et al., 2006c; Wang et al., 2006). Whereas Numb functions to antagonize Notch, Brat translationally represses the expression of Notch downstream effector genes (Haenfler et al., 2012; Komori et al., 2018; Landskron et al., 2018). In the type I lineage, simultaneous loss of *numb* and *brat* function leads to ectopic activation of Notch signaling in GMCs and the formation of supernumerary type I neuroblasts (Abdusselamoglu et al., 2019). In the type II lineage, mutation in *numb* or *brat* alone results in ectopic Notch activity in immature INPs and supernumerary type II neuroblasts (Bello et al., 2008; Boone and Doe, 2008; Bowman et al., 2008). Thus, mechanisms controlling the precise localization and segregation of Numb and Brat play a key role in regulating Notch-mediated binary cell fate decisions during asymmetric cell division.

Polarization of the cell cortex during asymmetric cell division provides a robust mechanism for segregating determinants to one of the two progeny, allowing one cell to rapidly acquire a new identity. The Par complex, consisting of Par-6 and atypical Protein Kinase C (aPKC), localizes to a crescent at the apical cortex of mitotic neuroblasts (Rolls et al., 2003). Mechanistic studies based on analyses of a few select aPKC downstream proteins, including Mira and Par-1, led to a “localization by exclusion” model in which aPKC negatively regulates cortical localization of its target proteins (Atwood and Prehoda, 2009; Bailey and Prehoda, 2015; Suzuki et al., 2004). Because phosphorylation by aPKC dissociates Mira from the apical cortex into the cytoplasm, Mira and its cargo protein Brat accumulate only at the basal cortex of mitotic neuroblasts and asymmetrically segregate into GMCs and immature INPs. Previous studies have suggested that aPKC regulates asymmetric Numb localization via an identical “localization by exclusion” mechanism (Smith et al., 2007; Wirtz-Peitz et al., 2008). Numb can be phosphorylated by aPKC at five evolutionarily conserved sites *in vitro*. Whereas Numb transgenic protein that is phosphomimetic at two of these sites asymmetrically localizes and segregates in mitotic neuroblasts, the non-phosphorylatable form of Numb transgenic protein remains uniformly localized in the neuroblast cortex (Haenfler et al., 2012). These results appear to contradict to the proposed “localization by exclusion” model and suggest that phosphorylation by aPKC promotes Numb localization to the basal cortex of mitotic neuroblasts and then to GMCs and immature INPs. However, direct evidence demonstrating that aPKC kinase activity directs basal Numb localization and segregation remains unavailable.

Here, we have identified Cullin 3 (Cul3) as a novel regulator of aPKC-directed basal localization and segregation of Numb during asymmetric neuroblast division. We have demonstrated that a *numb* hypomorphic allele drastically reduces Numb protein levels and leads to supernumerary neuroblasts when GMCs and immature INPs fail to downregulate Notch signaling and differentiate. In *cul3*-mutant brains, broader cortical localization of Numb in mitotic neuroblasts decreases the levels of Numb segregated into GMCs and immature INPs, leading to the formation of supernumerary neuroblasts. Surprisingly, increased aPKC kinase activity in *numb-* or *cul3*-mutant brains concentrates residual Numb protein at the basal cortex of mitotic neuroblasts and suppresses supernumerary neuroblast formation. By contrast, decreased aPKC function in either the *numb* or *cul3* genetic background randomizes Numb segregation and drastically enhances supernumerary neuroblast formation. We propose that Cul3 functions through aPKC-directed basal localization of Numb to regulate Notch-mediated binary cell fate decisions during asymmetric neuroblast division.

## Results

### Apical aPKC kinase activity concentrates Numb at the basal cortex of mitotic neuroblasts

To investigate the mechanisms controlling the spatial distribution of Numb during asymmetric neuroblast division, we generated a sensitized genetic background allowing for the detection of subtle changes in endogenous Numb localization and activity. Whereas *numb*-null animals die during embryogenesis, animals carrying the *numb*^*NP2301*^ allele in trans with a *numb*-null allele survive beyond larval development (Janssens et al., 2014; Komori et al., 2018). Because the *numb*^*NP2301*^ allele is induced by a transposable P-element inserted in the 5’ regulatory region of *numb*, we imprecisely excised the P-element to generate small deletions of the genomic region flanking the insertion site (Supplemental figure 1A). Similar to the *numb*^*NP2301*^ allele, animals carrying these new *numb*^*ex*^ alleles in trans with a *numb*-null allele also survived beyond larval development. We used the combination of the *numb*^*ex112*^ allele over a *numb*-null allele (abbreviated as *numb*^*hypo*^) for the remainder of this study.

**Figure 1.**
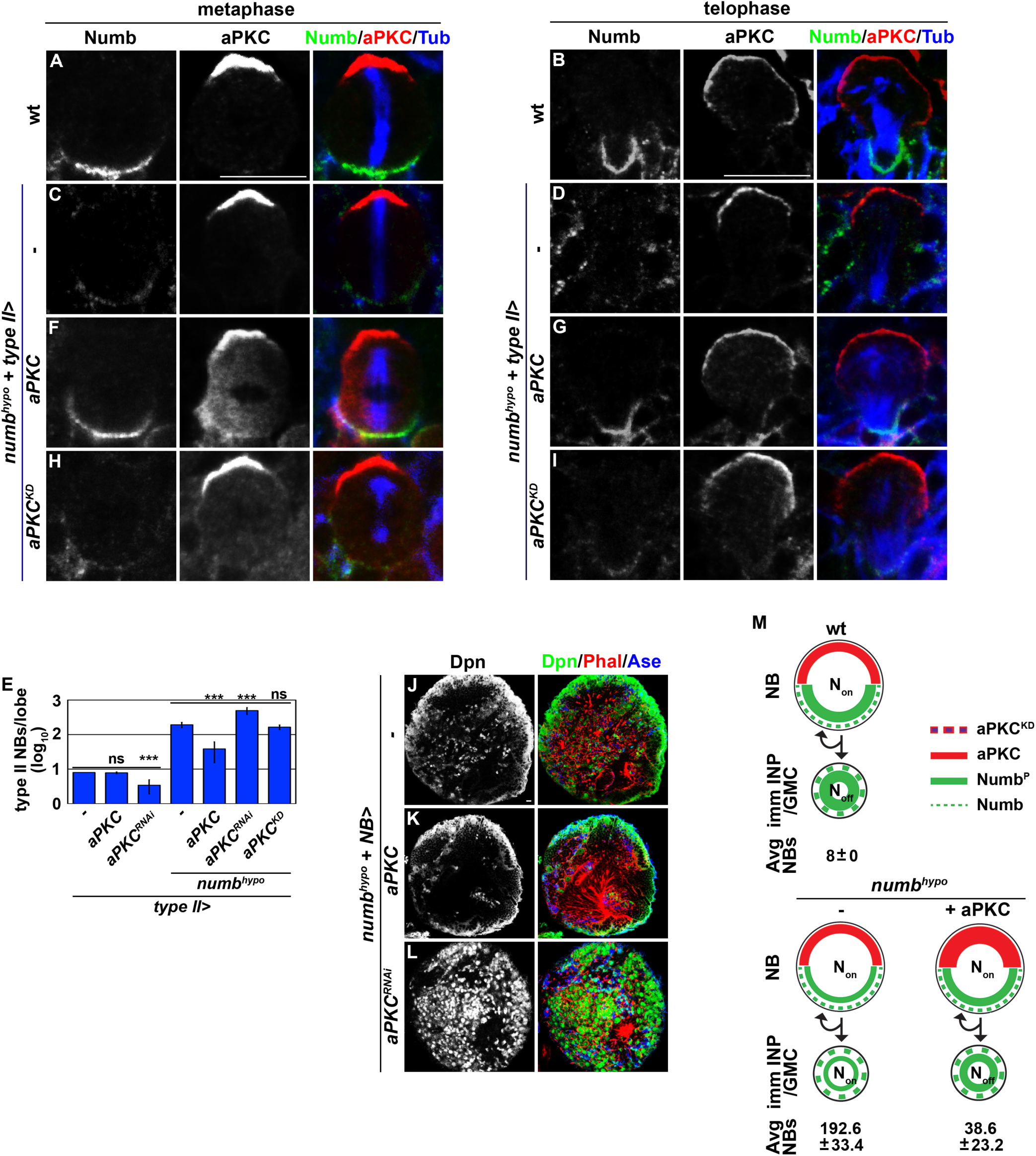
aPKC kinase activity concentrates Numb at the basal cortex of mitotic neuroblasts. (A-D) Metaphase and telophase *numb*^*hypo*^ (*numb*^*15/Ex112*^) neuroblasts showed much lower levels of Numb expression than wild-type neuroblasts. (E) The total type II neuroblasts are shown per brain lobe in wild-type or *numb*^*hypo*^ larvae overexpressing the indicated transgenes in type II neuroblasts. (F-I) Overexpressing wild-type aPKC but not kinase-dead aPKC (aPKC<u>^KD^</u>) restored basal enrichment of Numb in mitotic *numb*^*hypo*^ type II neuroblasts. (J-M) Increasing the expression of wild-type aPKC in *numb*^*hypo*^ brains suppressed the supernumerary neuroblast phenotype, whereas decreasing the expression of endogenous aPKC enhanced the phenotype. Overexpressing aPKC^KD^ in *numb*^*hypo*^ brains had no effect on the supernumerary neuroblast phenotype. (M) The diagram illustrates the aPKC and Numb localization patterns in neuroblasts of the indicated genotype, and presents a summary of total type II neuroblasts per lobe in these brains. NB: neuroblast. Type II NB>: type II neuroblast Gal4 driver (*Wor-Gal4,Ase-Gal80*). “-” indicates no transgene. Bar graphs are represented as mean ± standard deviation. P-values: *** <0.005. ns: not significant. Scale bar, 10 μm.

In wild-type brains, Numb asymmetrically accumulated in a basal crescent in metaphase type II neuroblasts and exclusively segregated into future immature INPs (Figures 1A,B). By contrast, *numb*^*hypo*^ brains showed significantly less Numb accumulation in the basal cortex of metaphase neuroblasts and segregated into immature INPs (Figures 1C,D). While wild-type brain lobes always contained eight type II neuroblasts, *numb*^*hypo*^ brain lobes contained approximately 250 supernumerary type II neuroblasts (Figure 1E). Thus, *numb*^*hypo*^ brains showed decreased Numb levels in immature INPs and contained supernumerary neuroblasts. Consistent with these findings, restoring Numb expression or reducing Notch-activated gene transcription by overexpressing a dominant-negative form of Mastermind (Man^DN^) in *numb*^*hypo*^ brains rescued the supernumerary neuroblast phenotype (Supplementary figures 1B-I). These data confirm that the decreased levels of Numb segregated into immature INPs in *numb*^*hypo*^ brains are insufficient to prevent ectopic Notch signaling and supernumerary neuroblast formation. We conclude that the *numb*^*hypo*^ genetic background provides a sensitized *in vivo* model for investigating the mechanistic control of Numb localization during asymmetric neuroblast division.

We overexpressed wild-type aPKC in *numb*^*hypo*^ brains to investigate the role of aPKC in regulating asymmetric Numb localization in mitotic neuroblasts. If the cortical exclusion model is correct, the increased aPKC levels should further delocalize Numb and enhance the supernumerary neuroblast phenotype. Indeed, we found that aPKC overexpression in both wild-type and *numb*^*hypo*^ brains expands cortical accumulation of aPKC beyond the normal apical domain, as expected (Figures 1A-D). Surprisingly, elevated cortical aPKC activity in *numb*^*hypo*^ brains <u>increased</u> basal Numb levels (Figures 1F,G). These data are inconsistent with the “cortical exclusion” model and instead show that aPKC can positively promote Numb basal localization. By contrast, overexpression of kinase-dead aPKC (aPKC^KD^) did not increase basal localization of Numb (Figures 1H,I). These data are consistent with our previous finding that the phosphomimetic but not the non-phosphorylatable form of transgenic Numb protein asymmetrically localizes to the basal cortex of mitotic neuroblasts (Haenfler et al., 2012). Thus, increasing wild-type aPKC kinase activity in *numb*^*hypo*^ brains increases Numb levels at the basal cortex of mitotic neuroblasts and Numb levels segregated into immature INPs. We conclude that aPKC can promote Numb basal localization.

We next determined the functional consequence of aPKC-induced basal localization in *numb*^*hypo*^ brains. *numb*^*hypo*^ brains contained approximately 250 supernumerary type II neuroblasts per lobe, compared with eight in wild-type brains (Figures 1E,J). Overexpressing wild-type aPKC in *numb*^*hypo*^ brains significantly suppressed the supernumerary neuroblast phenotype (Figures 1E,K). By contrast, knocking down *aPKC* function in *numb*^*hypo*^ brains drastically enhanced the supernumerary neuroblast phenotype (Figures 1E,L). Importantly, overexpressing aPKC^KD^ in *numb*^*hypo*^ brains had no effect on the supernumerary neuroblast phenotype (Figure 1E). Thus, increased wild-type aPKC activity in *numb*^*hypo*^ brains suppressed the reversion of immature INPs into supernumerary neuroblasts. Together, these data strongly support a model in which apical aPKC kinase activity positively promotes basal Numb localization in mitotic type II neuroblasts, allowing Numb to asymmetrically segregate into immature INPs and inhibit Notch signaling (Figure 1M).

### *cul3* is required for differentiation in neuroblast progeny

As aPKC-induced Numb basal localization is a new pathway, additional components in the pathway are not yet known. We reasoned that reducing the function of genes essential for aPKC-directed basal localization of Numb should enhance the supernumerary neuroblast phenotype in *numb*^*hypo*^ brains. We screened all lethal mutations induced by transposable P-element insertions on the second chromosome of the fly genome for dominant enhancers of the supernumerary type II neuroblast phenotype in *numb*^*hypo*^ brains. We found that heterozygosity of the *cul3*^*06430*^ allele enhanced the supernumerary neuroblast phenotype in *numb*^*hypo*^ brains (Figure 2A). Similarly, knocking down *cul3* function in *numb*^*hypo*^ brains enhanced the supernumerary neuroblast phenotype (Figure 2B). These results strongly suggest that reducing *cul3* function increases the reversion of immature INPs to supernumerary neuroblasts in *numb*^*hypo*^ brains. To determine whether loss of *cul3* function alone leads to supernumerary neuroblast formation, we examined the cell identity of *cul3*-mutant neuroblast clones marked by green fluorescent protein (GFP) expression. Control type I or type II neuroblast clones contained one neuroblast per clone at all time points examined, as expected (Figures 2C,E,F,H). By contrast, *cul3*-mutant neuroblast clones showed a progressively increased number of neuroblasts as the animals aged, indicating that some GMCs and immature INPs had reverted to neuroblasts (Figures 2D,E,G,H). Thus, we conclude that Cul3 promotes differentiation in GMCs and immature INPs.

**Figure 2.**
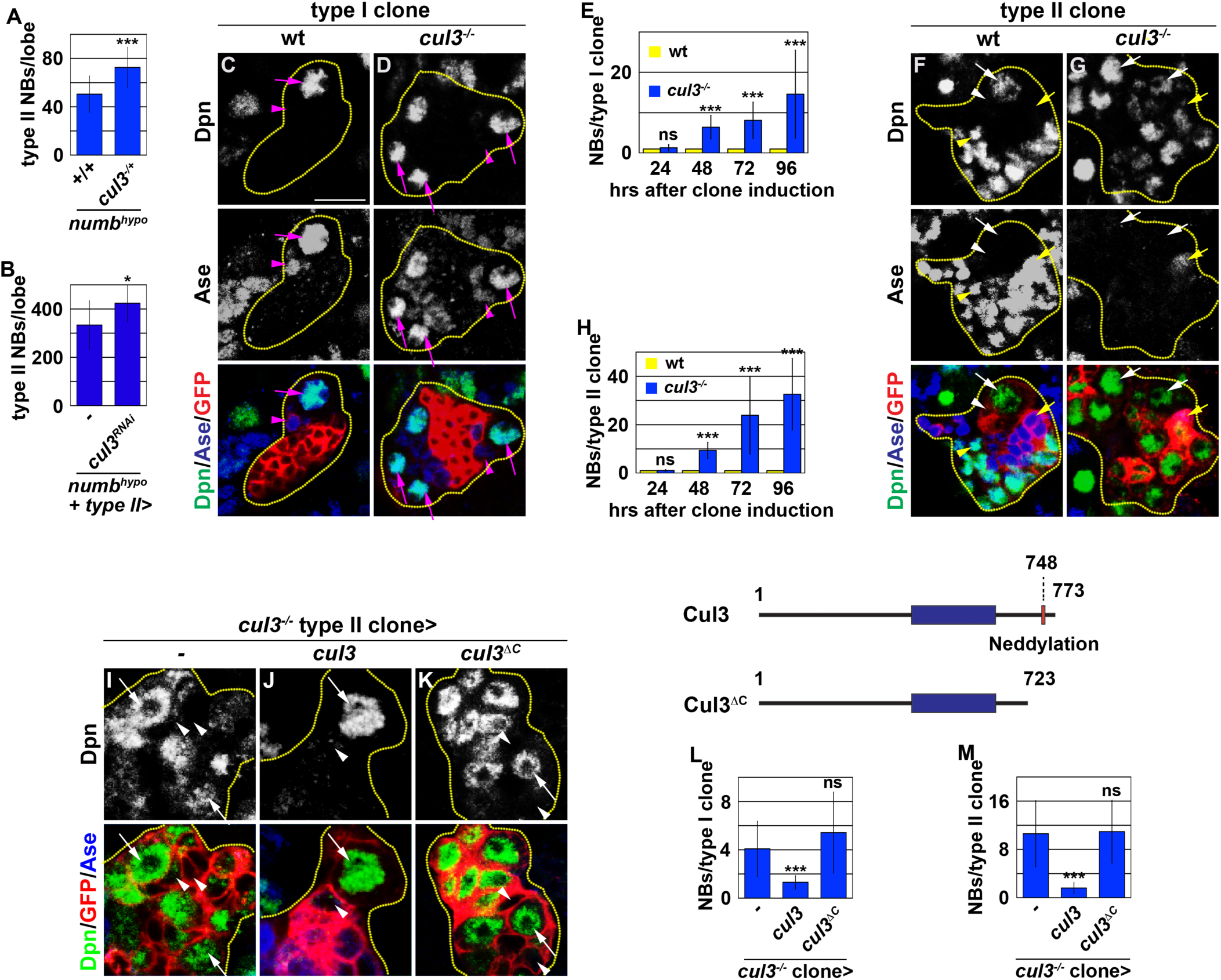
*cul3* promotes differentiation in neuroblast progeny. (A-B) A 50% reduction of *cul3* gene dosage or knockdown of *cul3* function by RNAi enhanced the supernumerary neuroblast phenotype in *numb*^*hypo*^ brains. (C-D) A GFP-marked wild-type type I neuroblast mosaic clone always maintained a single neuroblast, whereas a *cul3* homozygous mutant clone contained multiple neuroblasts. (E) A time-course analysis of total neuroblasts per wild-type or *cul3*-mutant type I neuroblast clone. (F-G) A wild-type type II neuroblast mosaic clone always maintained a single neuroblast, whereas a *cul3*-mutant clone contained multiple neuroblasts. (H) A time-course analysis of total neuroblasts per wild-type or *cul3*-mutant type II neuroblast clone. (I-K) Overexpressing wild-type Cul3 but not Cul3^ΔC^ rescued the supernumerary neuroblast phenotype in *cul3-*mutant type II neuroblast clones. (L-M) The average number of neuroblasts per *cul3*-mutant type I or type II neuroblast clone with or without overexpression of the indicated *UAS* transgene. Neuroblast clones are marked by GFP expression and are outlined in yellow dotted lines. The *cul3*^*06430*^ allele was used in these experiments. NB: neuroblast. Type II NB>: type II neuroblast Gal4 driver (*Wor-Gal4,Ase-Gal80*). “-” indicates no transgene. Magenta arrow indicates type I neuroblast (Dpn^+^Ase^+^). Magenta arrowhead indicates GMC (Dpn^-^Ase^+^). White arrow indicates type II neuroblast (Dpn^+^Ase^-^). White arrowhead indicates Ase^-^ immature INP (Dpn^-^Ase^-^). Yellow arrow indicates Ase^+^ immature INP (Dpn^-^Ase^+^). Bar graphs are represented as mean ± standard deviation. P-values: * <0.5, *** <0.005. ns: not significant. Scale bar, 10 μm.

Conjugation of the ubiquitin-like molecule Nedd8 to the C-terminus of Cullin-family proteins, including Cul3, promotes ubiquitinylation of their target proteins (Lydeard et al., 2013; Sarikas et al., 2011). We investigated whether Cul3 functions through its ubiquitin ligase activity to promote differentiation. Overexpressing wild-type Cul3 in *cul3*-mutant clones rescued the supernumerary neuroblast phenotype and restored differentiation (Figures 2I,J,L,M). By contrast, overexpressing a *UAS-cul3*^*ΔC*^ transgene that encodes transgenic Cul3 protein lacking the Neddylation motif in the C-terminus in *cul3*-mutant clones did not rescue the supernumerary neuroblast phenotype (Figures 2K-M). Thus, the ubiquitin ligase activity of Cul3 is necessary to promote differentiation in GMCs and immature INPs.

### Cul3 promotes differentiation in neuroblast progeny by downregulating Notch signaling

Notch functions through transcription factors encoded by *deadpan* (*dpn*) and *Enhancer of Split mγ* [*E(spl)mγ*] to maintain neuroblasts in an undifferentiated state, and downregulation of Notch signaling is required for neuroblast progeny to differentiate into GMCs and immature INPs. We evaluated the expression of these Notch target genes to determine whether Cul3 promotes differentiation in GMCs and immature INPs by downregulating Notch signaling. In wild-type clones, we detected Dpn and E(spl)mγ in all type I and type II neuroblasts, but not in GMCs in the type I neuroblast lineage or in immature INPs in the type II neuroblast lineage (Figures 3A,C). In *cul3*-mutant clones, however, we detected Dpn and E(spl)mγ in both neuroblasts and their progeny (Figures 3B,D). These results support a model in which *cul3* promotes differentiation in GMCs and immature INPs by downregulating Notch signaling.

**Figure 3.**
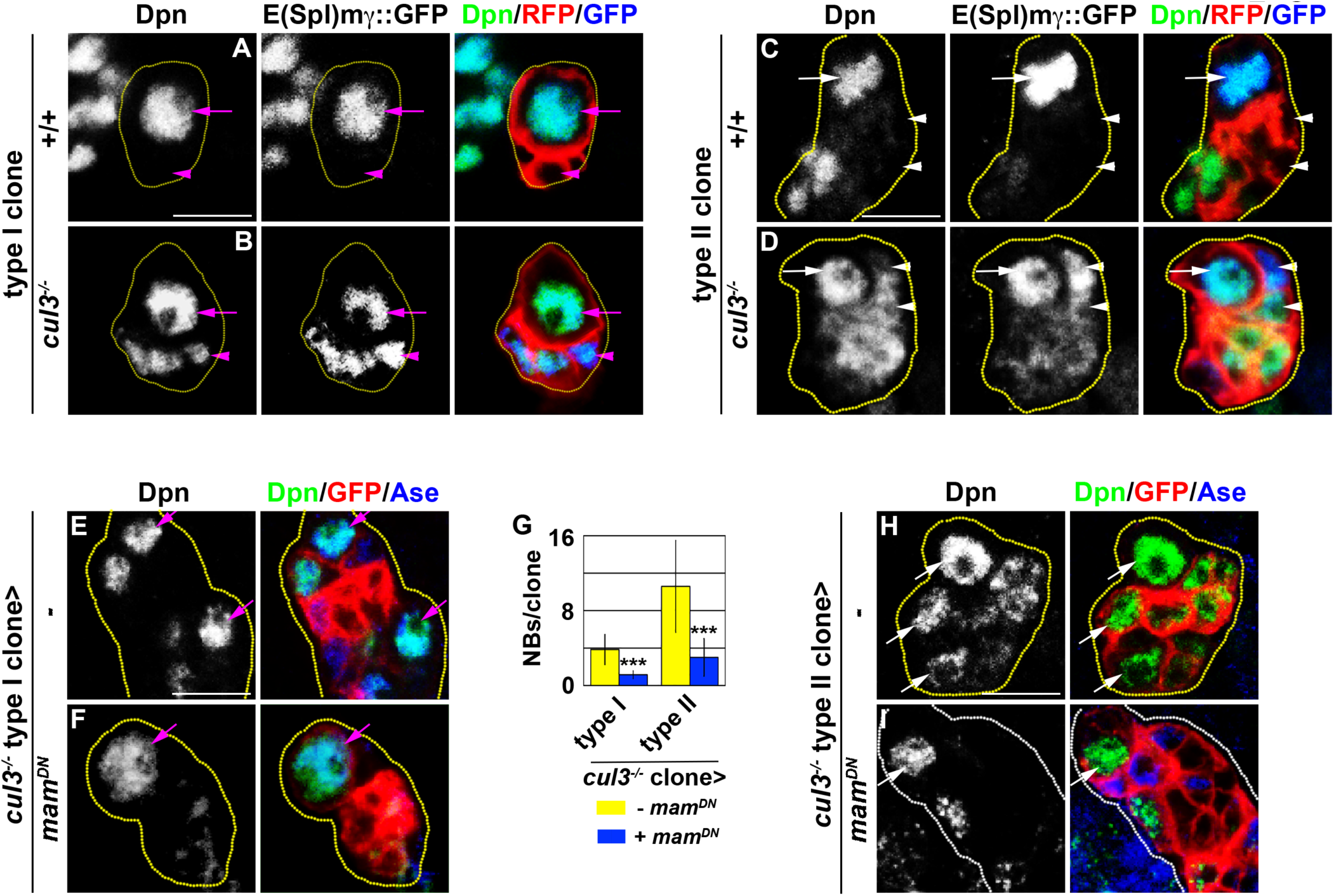
Cul3 suppresses transcriptional activation of Notch target genes in neuroblast progeny. (A-D) Notch target genes *dpn* and *E(spl)mγ* were not expressed in the progeny of wild-type neuroblasts, but were aberrantly expressed in the progeny of *cul3* homozygous mutant neuroblasts. Neuroblast clones are marked by RFP expression and are outlined in yellow dotted lines. (E-I) Inhibiting transcriptional activation of Notch target genes by overexpressing Mam^DN^ suppressed supernumerary neuroblast formation in *cul3*-mutant type I and type II neuroblast clones. The average number of neuroblasts per *cul3-*mutant type I or type II neuroblast clone with or without Mam^DN^ overexpression is shown in G. Neuroblast clones are marked by GFP expression and are outlined in yellow dotted lines. The *cul3*^*06430*^ allele was used in these experiments. Magenta arrow indicates type I neuroblast (Dpn^+^Ase^+^). Magenta arrowhead indicates GMC (Dpn^-^Ase^+^). White arrow indicates type II neuroblast (Dpn^+^Ase^-^). White arrowhead indicates Ase^-^ immature INP (Dpn^-^Ase^-^). Bar graphs are represented as mean ± standard deviation. P-values: *** <0.005. Scale bar, 10 μm.

We overexpressed a *UAS-mam*^*DN*^ transgene in *cul3*-mutant neuroblast clones to determine whether Notch-activated transcription of target genes contributes to supernumerary neuroblast formation. Mam^DN^ overexpression did not lead to premature neuroblast differentiation in wild-type clones, indicative of a mild reduction in Notch-activated target gene transcription (data not presented). By contrast, Mam^DN^ overexpression in *cul3*-mutant clones strongly suppressed the supernumerary neuroblast phenotype (Figures 3E-I). Thus, aberrant Notch-activated *dpn* and *E(spl)mγ* transcription in *cul3*-mutant GMCs and immature INPs leads to supernumerary neuroblast formation. We conclude that Cul3 promotes differentiation in GMCs and immature INPs by downregulating Notch signaling.

### Cul3 functions through Numb to promote differentiation in neuroblast progeny

Asymmetric segregation of Brat and Numb is required to downregulate Notch signaling in GMCs and immature INPs. Thus, reduced Numb or Brat activity in *cul3*-mutant GMCs and immature INPs could lead to supernumerary neuroblast formation. To distinguish these two possibilities, we overexpressed *brat* or *numb* in *cul3*-mutant type I and II neuroblast clones. Brat overexpression in *cul3*-mutant clones failed to suppress the supernumerary neuroblast phenotype (Figures 4A,B,D-F,H). Thus, it is unlikely that reduced Brat activity in *cul3*-mutant GMCs and immature INPs led to supernumerary neuroblast formation. Under identical conditions, Numb overexpression in *cul3*-mutant clones strongly suppressed the supernumerary neuroblast phenotype (Figures 4C,D,G,H). As a control, we confirmed that Numb overexpression in wild-type clones did not induce premature neuroblast differentiation (Figures 4D,H). Taken together, our results indicate that reduced Numb activity in *cul3*-mutant GMCs and immature INPs causes their reversion to supernumerary neuroblasts, and strongly suggest that Cul3 functions through Numb to downregulate Notch signaling.

**Figure 4.**
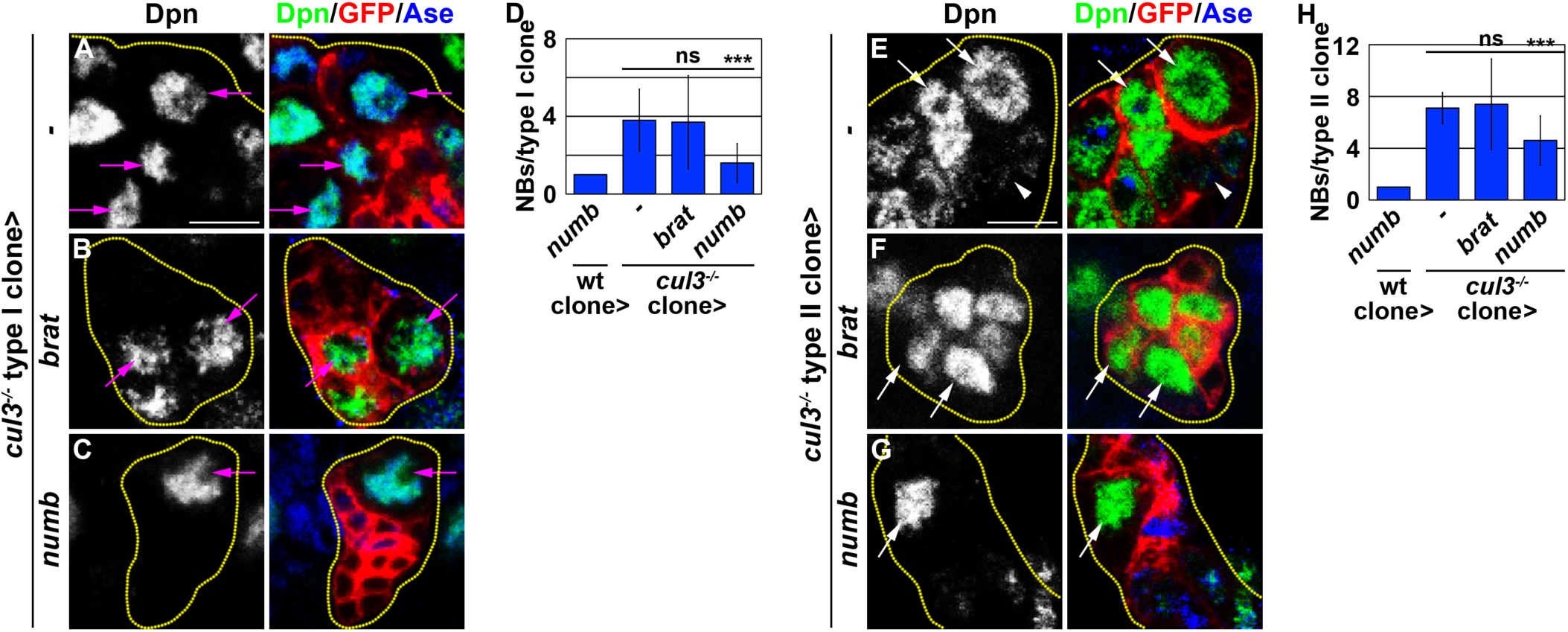
Numb overexpression suppressed supernumerary neuroblast formation in *cul3*-mutant neuroblast clones. (A-H) Overexpressing *numb* but not *brat* suppressed supernumerary neuroblast formation and restored differentiation in *cul3* homozygous mutant type I and type II neuroblast clones. The average numbers of neuroblasts per *cul3-*mutant type I or type II neuroblast clone with or without overexpression of indicated *UAS* transgenes are shown in D and H. NB: neuroblast. “-” indicates no transgene. Magenta arrow indicates type I neuroblast (Dpn^+^Ase^+^). White arrow indicates type II neuroblast (Dpn^+^Ase^-^). White arrowhead indicates Ase^-^ immature INP (Dpn^-^Ase^-^). Bar graphs are represented as mean ± standard deviation. The *cul3*^*06430*^ allele was used in these experiments. P-values: *** <0.005. Scale bar, 10 μm.

### *cul3* promotes asymmetric localization of Par-6/aPKC and Numb in mitotic neuroblasts

aPKC-induced Numb basal localization during asymmetric neuroblast division allows Numb to exclusively segregate into GMCs and immature INPs, where it downregulates Notch activation and promotes differentiation. Thus, Cul3 might downregulate Notch signaling by functioning in the aPKC-induced Numb basal localization pathway. We tested this hypothesis by first examining the localization pattern of Par proteins in mitotic type II neuroblasts. In wild-type neuroblasts, Par-6 and aPKC localized to an apical crescent, whereas loss of *cul3* function broadened the localization of Par-6 and aPKC to the basal-lateral cell cortex (Figures 5A,B,G). By contrast, the fly homolog of Par-3, Bazooka (Baz), remained localized to an apical crescent in mitotic wild-type and *cul3*-mutant neuroblasts (Figures 5C,D,G). Thus, *cul3* is required to maintain Par-6/PKC at the apical cortex of mitotic neuroblasts. Because binding to Par-6 activates the kinase activity of aPKC, co-localization of Par-6 and aPKC beyond the apical cortex of mitotic *cul3*-mutant neuroblasts suggests broader cortical aPKC kinase activity. We explored this possibility by examining the localization pattern of Mira, which delocalizes from the cortex into the cytoplasm upon phosphorylation by aPKC. Mira formed a basal crescent in metaphase neuroblasts among wild-type mitotic neuroblasts, but became uniformly localized in the cytoplasm of *cul3*-mutant neuroblasts (Figures 5C,D,G). Thus, Cul3 is a novel regulator of Par-6/aPKC apical localization in mitotic neuroblasts, with *cul3*-mutant neuroblasts displaying broader cortical aPKC activity.

**Figure 5.**
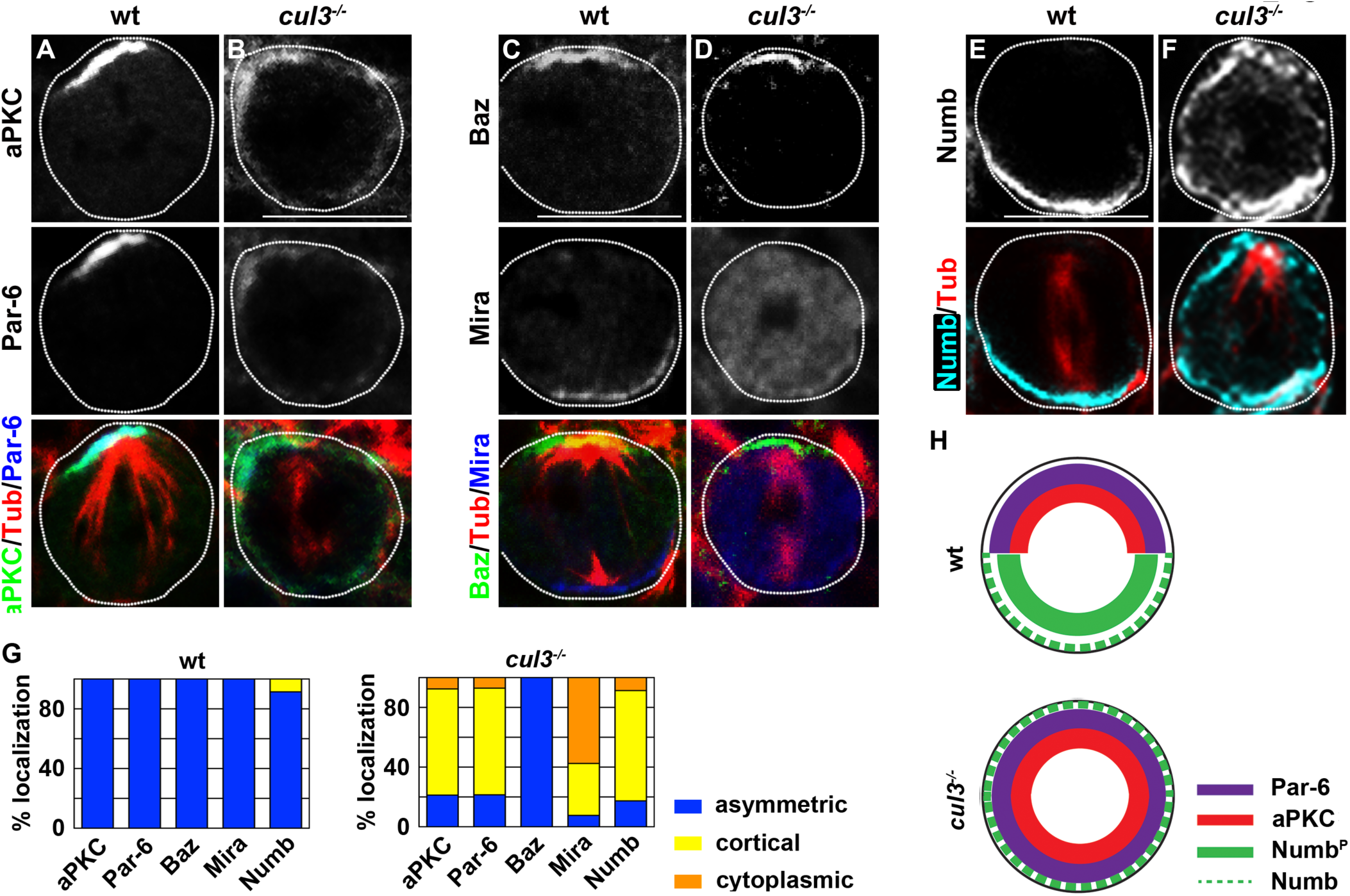
Cul3 is required for asymmetric localization of Par-6/aPKC and Numb in mitotic neuroblasts. (A-B) aPKC and Par-6 were localized in an apical crescent in wild-type metaphase neuroblasts, but ectopically localized in the cortex of *cul3*-mutant neuroblasts. (C-D) Baz was localized in an apical crescent in both wild-type and *cul3*-mutant neuroblasts. Mira was localized in a basal crescent in wild-type neuroblasts but mislocalized in the cytoplasm of *cul3*-mutant neuroblasts. (E-F) Numb was localized in an apical crescent in wild-type metaphase neuroblasts, but ectopically localized in the cortex of *cul3*-mutant neuroblasts. (G) The frequency of the localization patterns of the proteins examined above are shown for wild-type and *cul3*-mutant neuroblasts. (H) The diagram illustrates the localization pattern of aPKC, Par-6, and Numb in wild-type or *cul3*-mutant neuroblasts. Neuroblast cortex is outlined in white dotted lines. The *cul3*^*06430*^ allele was used in these experiments. Scale bar, 10 μm.

Surprisingly, *cul3*-mutant neuroblasts exhibited uniform cortical localization of Numb (Figures 5E-G). This result appears contradictory to our finding that cortical aPKC kinase activity positively promotes basal localization of Numb in mitotic neuroblasts. We previously demonstrated that the non-phosphorylatable form of transgenic Numb protein uniformly localizes throughout the cortex of mitotic neuroblasts (Haenfler et al., 2012). Thus, Numb likely remained unphosphorylated by aPKC in mitotic *cul3*-mutant neuroblasts despite broader cortical aPKC kinase activity. These data support our hypothesis that Cul3 functions through aPKC-directed basal localization of Numb to prevent ectopic activation of Notch in GMCs and immature INPs and suggest that Cul3 promotes phosphorylation of Numb by aPKC.

### Cul3 is required for aPKC-directed asymmetric localization and segregation of Numb

As a key open question, it remains unclear why Numb fails to localize to the basal cortex of *cul3*-mutant neuroblasts despite broader cortical aPKC kinase activity; this result is inconsistent with our observation that elevated aPKC levels promote Numb basal localization (Figure 1). One possibility is that Cul3 might promote aPKC phosphorylation of Numb by facilitating efficient substrate recognition, with the majority of Numb in the cortex of *cul3*-mutant neuroblasts being unable to be efficiently phosphorylated by aPKC and thus unable to target to the basal cortex. This hypothesis predicts that decreased aPKC function in mitotic *cul3*-mutant neuroblasts should further decrease basal Numb localization, whereas increased aPKC function should restore basal enrichment of Numb. We observed uniform cortical Numb localization in 30-45% of mitotic *aPKC* or *cul3* single-mutant neuroblasts (Figures 6A-C; Supplemental figure 2)

**Figure 6.**
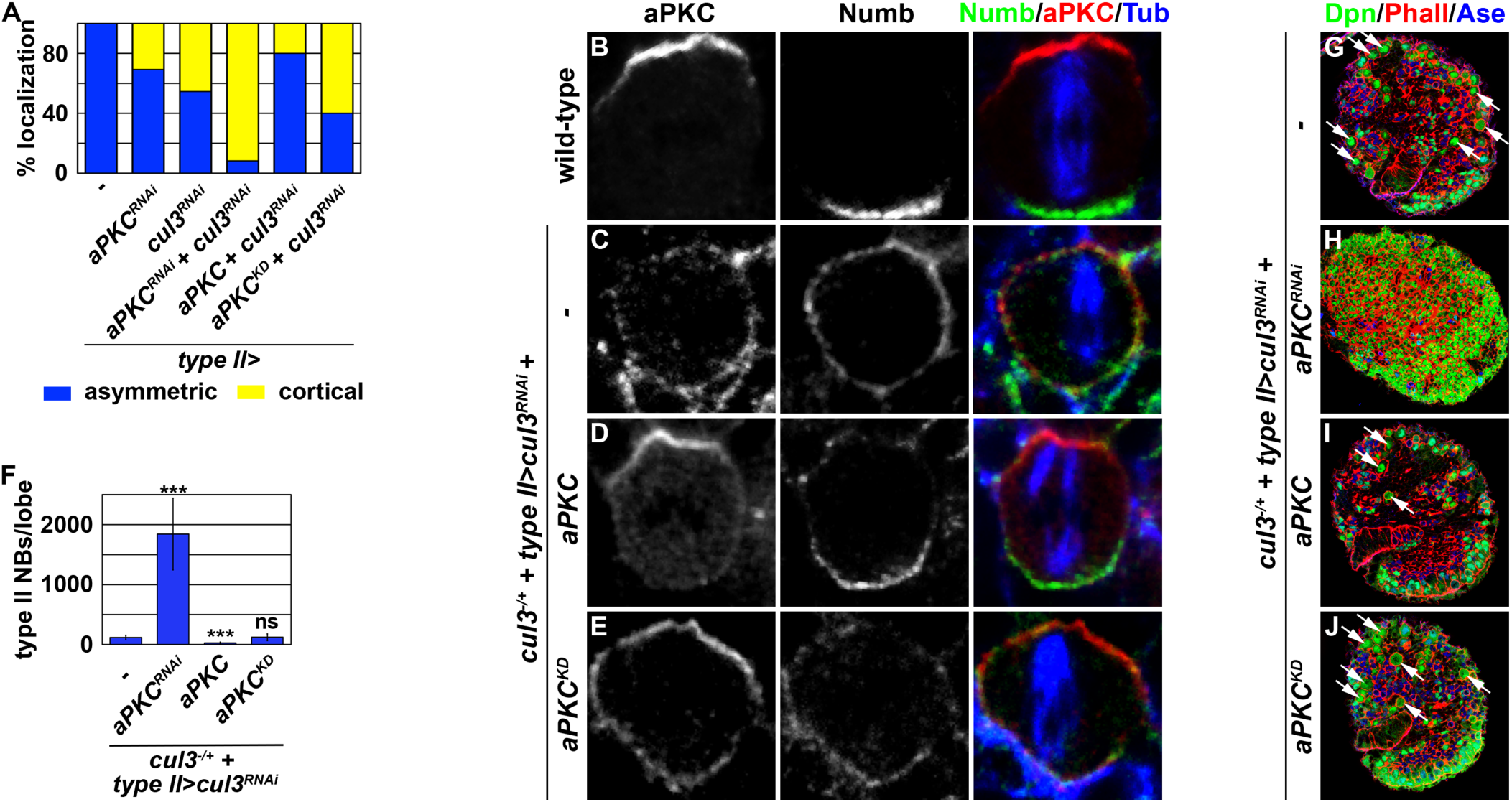
aPKC overexpression restores asymmetric segregation of Numb and suppressed supernumerary neuroblast formation in *cul3*-mutant brains. (A) The frequency of Numb localization patterns in wild-type neuroblasts or neuroblasts with indicated genes knocked down by RNAi. (B-E) aPKC and Numb were ectopically localized in the cortex of metaphase *cul3*-mutant neuroblasts. Overexpressing wild-type aPKC but not kinase-dead aPKC (aPKC^KD^) restored basal enrichment of Numb in metaphase *cul3*-mutant neuroblasts. (F) The average number of type II neuroblasts per brain lobe is shown for *cul3-*mutant brains with or without overexpression of the indicated *UAS* transgenes. (G-J) Knocking down *aPKC* function by RNAi enhanced supernumerary neuroblast formation in *cul3*-mutant brains. Overexpressing wild-type aPKC but not kinase-dead aPKC (aPKC^KD^) suppressed supernumerary neuroblast formation in *cul3*-mutant brains. White arrowhead indicates Ase^-^ immature INP (Dpn^-^Ase^-^). Bar graphs are represented as mean ± standard deviation. The *cul3*^*06430*^ allele was used in these experiments. P-values: *** <0.005. Scale bar, 10 μm.

). By contrast, 90% of mitotic *aPKC,cul3* double-mutant neuroblasts showed uniform cortical Numb localization (Figure 6A). Importantly, overexpressing wild-type aPKC but not aPKC^KD^ in mitotic *cul3*-mutant neuroblasts restored basal enrichment of Numb (Figures 6A,D,E). Thus, decreasing aPKC function in mitotic *cul3*-mutant neuroblasts further decreased basal Numb localization whereas increased aPKC function restored Numb basal localization.

Further randomization of cortical Numb localization in mitotic *cul3*-mutant neuroblasts when aPKC is decreased should reduce Numb levels segregated into immature INPs and enhance the supernumerary neuroblast phenotype. Indeed, knocking down *aPKC* function strongly enhanced the supernumerary neuroblast phenotype in *cul3*-mutant brains (Figures 6G-H). By contrast, the restoration of basal enrichment of Numb in mitotic *cul3*-mutant neuroblasts when aPKC function is increased should elevate Numb levels segregated into immature INPs and suppress the supernumerary neuroblast phenotype. Consistent with this prediction, overexpressing wild-type aPKC but not aPKC^KD^ strongly suppressed the supernumerary neuroblast phenotype in *cul3*-mutant brains (Figures 6F,I,J). These data demonstrate that increasing aPKC kinase activity in mitotic *cul3*-mutant neuroblasts can restore functional levels of Numb in the basal cortex, and thus in immature INPs, to allow for differentiation instead of reversion to supernumerary neuroblasts. We propose that Cul3 facilitates efficient phosphorylation of Numb by aPKC in mitotic neuroblasts, and thus exerts positive activity in the newly identified aPKC-induced Numb basal localization pathway.

## Discussion

Asymmetric segregation of Numb provides an evolutionarily conserved mechanism for eliciting Notch-mediated binary cell fate decisions following asymmetric cell division. The Par-6/aPKC complex plays a key role in asymmetrically localizing Numb, but the mechanisms by which Numb localizes to the opposite cell cortex from Par-6/aPKC remain poorly understood. In this study, we demonstrated that aPKC positively promotes basal Numb localization. In addition, we identified Cul3 as a positively acting component of this new aPKC-directed basal protein localization mechanism. We provide genetic and functional evidence supporting a model in which Cul3 promotes asymmetric sorting of Numb by facilitating efficient phosphorylation of Numb by aPKC (Figure 7). We propose that Cul3 functions through aPKC-directed asymmetric Numb localization and segregation to regulate Notch-mediated binary cell fate decision during asymmetric neuroblast division.

**Figure 7.**
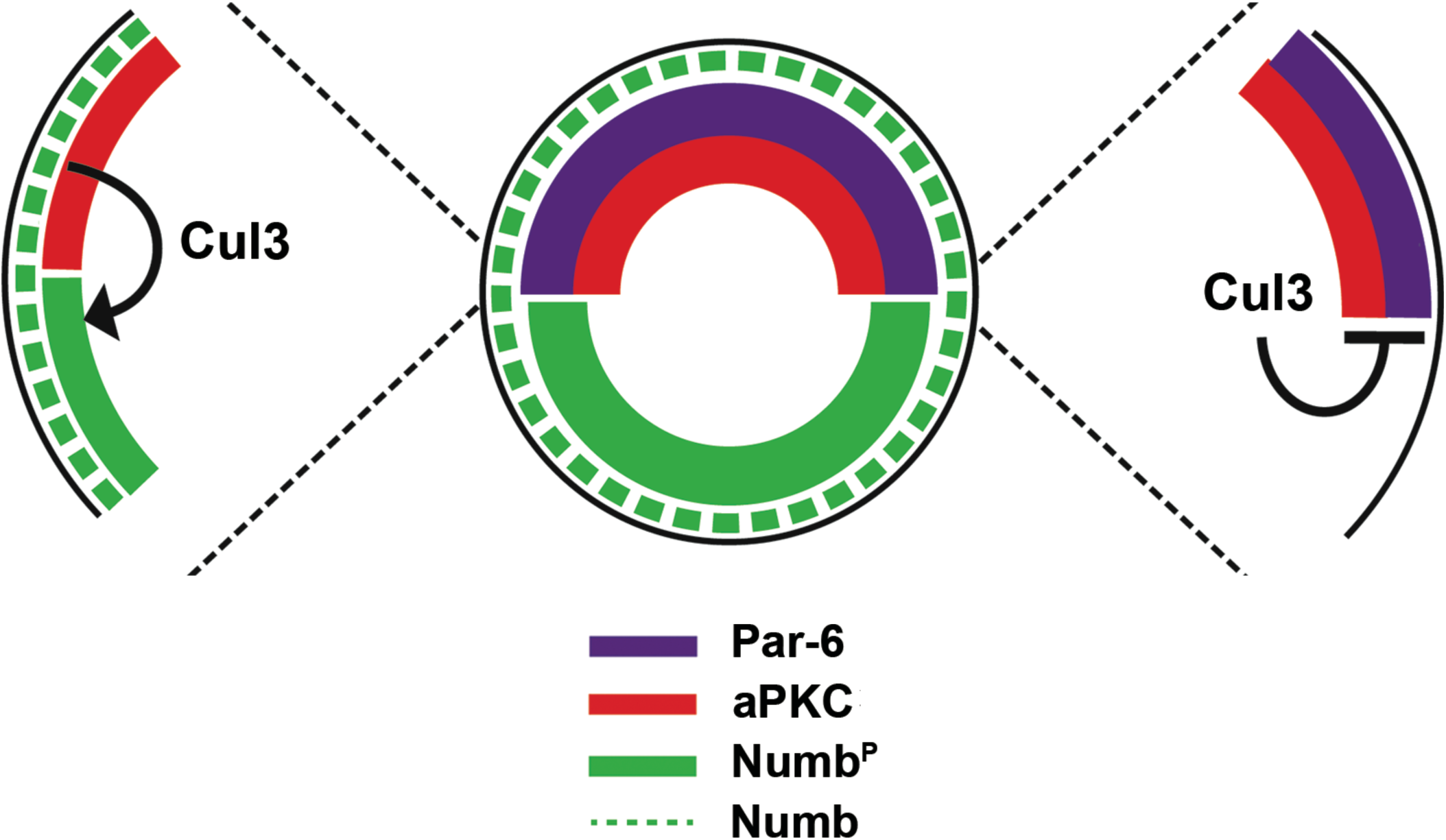
A model summarizing the functions of Cul3 during asymmetric neuroblast division. Cul3 is required for aPKC-directed localization of Numb in the basal cortex of mitotic neuroblasts, and is also required for maintaining the Par-6/aPKC complex at the apical cortex.

### aPKC-directed basal localization of Numb during asymmetric neuroblast division

Polarized localization of the Par-6/aPKC complex provides an evolutionarily conserved mechanism for asymmetrically localizing its downstream proteins in the cell cortex (Drummond and Prehoda, 2016; Lang and Munro, 2017). Mechanistic studies based on analyses of a few select aPKC downstream proteins, including Mira and Par-1, led to a “localization by exclusion” model (Atwood and Prehoda, 2009; Suzuki et al., 2004). This model postulates that phosphorylation by aPKC dissociates its downstream proteins from the cortex into the cytoplasm, leading to unphosphorylated proteins localizing to the cell cortex opposite to the aPKC cortical domain. The extent to which this “localization by exclusion” model is applicable to all polarized proteins remains unclear. More importantly, a cortical exclusion mechanism does not preclude additional parallel mechanisms. Indeed, we uncovered a parallel mechanism by which aPKC kinase activity directs additional downstream proteins to the basal cortex. This process could synergize with the cortical exclusion mechanism, leading to a robust and generalizable model for aPKC-regulated asymmetric protein localization in development and maintenance of homeostasis.

A sensitive experimental platform is essential for unraveling the mechanisms by which aPKC promotes asymmetric localization of downstream proteins. We leveraged *numb*^*hypo*^ brains, which provide a highly sensitized genetic background to screen for pathway components compared to wild-type or *numb-*null mutants (Figure 1). By using this experimental paradigm, our study provided genetic and functional evidence demonstrating that aPKC kinase activity positively promotes asymmetric Numb localization in mitotic neuroblasts. We found consistent findings using Numb^S2D^ transgenic proteins. When we replaced two conserved aPKC phosphorylation sites (S48 and S52) with aspartic acid, the Numb^S2D^ transgenic protein asymmetrically localized and segregated in mitotic neuroblasts; by contrast, when we replaced S48 and S52 with alanine, the protein symmetrically segregated (Haenfler et al., 2012). Thus, aPKC regulates asymmetric localization of some downstream proteins (Mira and Par-1) by cortical exclusion, and other proteins (Numb) by positively promoting basal localization.

The cellular machinery underlying the aPKC-Numb positive localization pathway appears to significantly differ from the “localization by exclusion” pathway components. For example, the phosphorylation state of the Numb S48 and S52 residues uniquely contributes to aPKC-directed basal localization of Numb during asymmetric neuroblast division while having no effect on Numb-mediated antagonism of Notch signaling (Haenfler et al., 2012). A key future experiment will be to use CRISPR genome engineering to generate *numb* alleles encoding Numb^S2D^ and Numb^S2A^ in order to confirm that the phosphorylation state of these two residues is indeed required for asymmetric Numb localization. Once confirmed, it will be important to identify proteins that uniquely translocate Numb^S2D^ to the basal cortex of mitotic neuroblasts. Findings from such experiments would provide insight into the functional differences between the regulation of Numb localization and the regulation of Notch signaling by Numb, and could be broadly applicable to many other cell types and model systems.

### Cul3 is a multifaceted regulator of cortical polarity and Notch-mediated binary cell fate decisions

Genetic studies in model organisms have led to the emerging concept that different Cullins evolved to fulfill specialized cellular functions (Sarikas et al., 2011). For example, the Cul3 ubiquitin ligase complex likely functions as a key regulator of tumor suppression and cell differentiation in mammals (Chen and Chen, 2016; Dubiel et al., 2018). In this study, we found that Cul3 is required for maintaining apical-basal cortical polarity and downregulating Notch signaling during asymmetric neuroblast division. Loss of apical-basal cortical polarity is a hallmark of tumorigenesis, whereas defects in the termination of Notch activity prevent the timely onset of differentiation in many stem cell lineages. Thus, our findings provide novel insights into how Cul3 might suppress tumorigenesis and promote differentiation.

Similar to *cul3*, the tumor-suppressor gene *lethal giant larvae* (*lgl*) regulates cell polarity and promotes differentiation during asymmetric neuroblast division (Lee et al., 2006b). While both *lgl* and *cul3*-mutant brains display supernumerary neuroblasts and ectopic cortical aPKC localization in mitotic neuroblasts, the underlying mechanisms appear to be different. In mitotic *lgl*-mutant neuroblasts, multiple apical polarity proteins, including Par6/aPKC, Baz, and Partner of inscutable (Pins), are mislocalized. In contrast, Baz and Pins remain localized in an apical crescent in mitotic *cul3*-mutant neuroblasts. Moreover, reducing *aPKC* function *suppresses* supernumerary neuroblast formation in *lgl*-mutant brains but *enhances* this phenotype in *cul3*-mutant brains. Finally, Lgl globally regulates the localization of most, if not all, apical polarity proteins during asymmetric neuroblast division, whereas Cul3 specifically regulates Par-6/aPKC apical localization (Figure 7). A key open question is how Cul3 restricts the Par-6/aPKC complex to the apical cortex. Mislocalization of Mira from the basal cortex to the cytoplasm in mitotic *cul3*-mutant neuroblasts indicates that the cortex exhibits ectopic aPKC kinase activity (Figure 5D). Binding of the Par-6/aPKC complex to Rho GTPase Cdc42 enables activation of cortical aPKC kinase activity (Rodriguez et al., 2017). Thus, it is plausible that increased cortical Cdc42 activity might contribute to ectopic cortical aPKC kinase activity in mitotic *cul3*-mutant neuroblasts. In mammals, Cul3 coordinates the early steps of cortical neurogenesis by reorganizing the actin cytoskeleton via Rho GTPases (Gladwyn-Ng et al., 2015; Pacary et al., 2013). We speculate that Cul3 might maintain the apical localization of the Par-6/aPKC complex by regulating Rho GTPase such as Cdc42 and actin cytoskeleton dynamics.

Failure to asymmetrically localize Numb in mitotic *cul3*-mutant neuroblasts despite exhibiting broader cortical aPKC kinase activity appears to contradict the model that aPKC positively directs basal localization of Numb. One possibility is that *cul3* loss of function perturbs a separate mechanism that functions in parallel with aPKC to localize Numb to the basal cortex of mitotic neuroblasts. Another possibility is that aPKC must be polarized to the apical cortex to promote basal Numb localization. Alternatively, Numb might not be efficiently phosphorylated by aPKC in mitotic *cul3*-mutant neuroblasts, and thus fails to undergo aPKC-directed basal localization. Our data favor the latter possibility because overexpressing wild-type but not kinase-dead aPKC restores asymmetric Numb localization in mitotic *cul3*-mutant neuroblasts and suppresses the supernumerary neuroblast phenotype in *cul3*-mutant brains. We propose that Cul3 promotes an efficient interaction between Numb and aPKC, and overexpressed aPKC compensates for a reduced aPKC-Numb interaction by increasing phosphorylation of Numb and concentrating it to the basal cortex of mitotic *cul3*-mutant neuroblasts. Because the E3 ubiquitin ligase activity of Cul3 is required to suppress supernumerary neuroblast formation, we speculate that the non-proteolytic function of Cul3 facilitates an efficient interaction between aPKC and Numb. More experiments are required to test this hypothesis and elucidate the regulation of substrate recognition by aPKC.

## Materials and Methods

### Fly genetics and transgenes

We used Oregon R as the wild-type control and the following transgenic and mutant strains: *Wor-Gal4* (Lee et al., 2006b), *brat*^*DG19310*^ (Xiao et al., 2012), *numb*^*15*^ (Berdnik et al., 2002), *Ase-Gal80* (Neumüller et al., 2011), *E(spl)mγ-GFP* (Almeida and Bray, 2005), *UAS-HA::aPKC, UAS-aPKC*^*K293W*^ (kindly provided by Dr. K. Prehoda), *UAS-brat-myc* (Komori et al., 2014), *UAS-numb-myc* (Haenfler et al., 2012), and *UAS-mam*^*DN*^ (Kumar and Moses, 2001). The following stocks were obtained from the Bloomington *Drosophila* Stock Center: *brat*^*11*^, *cul3*^*06430*^, *Elav-Gal4 (C155), FRT40A, hs-flp, UAS-aPKC*^*RNAi*^ (TRiP.HMS01320), *UAS-cul3*^*RNAi*^ (TRiP.HM05109), *Tub-Gal80, Tub-Gal80*^*ts*^, *UAS-CD4-tdTom, UAS-gft*^*FL*^, *UAS-gft*^*ΔC*^, *UAS-dcr2, UAS-mCD8::GFP, Tub-Gal80*, and *Tub-Gal80*^*ts*^.

### Clonal analyses

Lineage clones were induced following a previously published method (Lee and Luo, 2001). Briefly, larvae were incubated at 37°C for 90 minutes for heat shock. After heat shock, larvae were cultured at 25°C until dissection.

### Edu labeling

Larvae were cultured until they reached the second or third instar stage and were fed food containing 0.2 mM Edu for 3 hours. After 3 hours of Edu incorporation, some larvae were dissected for pulse experiments; larvae for the chase experiment were cultured in regular food for an extra 20 hours before dissection. After dissection, fixation and Edu labeling were performed following a previously published method (Daul et al., 2010).

### Immunofluorescence staining and antibodies

Larvae brains were dissected in phosphate-buffered saline (PBS) and fixed in 100 mM Pipes (pH 6.9), 1 mM EGTA, 0.3% Triton X-100, and 1 mM MgSO_4_ containing 4% formaldehyde for 23 minutes. Fixed brain samples were washed with PBS and 0.3% Triton X-100 (i.e., PBST). After removing the fix solution, samples were incubated with primary antibodies for 3 hours at room temperature. Three hours later, samples were washed with PBST, and then incubated with secondary antibodies overnight at 4°C. On the next day, samples were washed with PBST and equilibrated in ProLong Gold antifade mount (ThermoFisher Scientific). Antibodies used in this study include chicken anti-GFP (1:2,000; Aves Labs), mouse anti-tub (1:2,000; Sigma), guinea pig anti-Baz (1:100) (Siller et al., 2006), guinea pig anti-Numb (1:500) (O’Connor-Giles and Skeath, 2003), rabbit anti-aPKC (1:1,000; Sigma), rabbit anti-Ase (1:400) (Weng et al., 2010), rabbit anti-RFP (1:200; Rockland), rat anti-Dpn (1:2) (Lee et al., 2006b), rat anti-Mira (1:500) (Lee et al., 2006b), and rat anti-Par-6 (1:600) (Rolls et al., 2003). Secondary antibodies were from Jackson ImmunoResearch Inc. We used rhodamine phalloidin (ThermoFisher Scientific) to visualize cortical actin. Confocal images were acquired on a Leica SP5 scanning confocal microscope (Leica Microsystems Inc). More than ten brains per genotype were used to obtain data for each experiment.

### Quantification and Statistical Analysis

The standard deviation among samples is indicated by error bars. All statistical analyses were performed using a two-tailed Student’s T-test. P-values < 0.05, <0.005, and <0.0005 are indicated by (*), (**), and (***), respectively, in the figures.

## Acknowledgments

We thank Drs. K Prehoda and A. Wodarz for providing reagents. We thank the Bloomington *Drosophila* Stock Center for fly stocks. We thank Science Editors Network for editing the manuscript. We thank Ms. Alina Moroz, Ms. Caroline Poirier, and Ms. Julia Derringer for technical assistance, and Cyrina Ostgaard for graphic illustration. This work was supported by a National Institutes of Health grant (R01NS107496) to C.-Y.L.

## Author Contributions

Conceptualization: H. K., N. R.-Q., and C.-Y. L.

Formal analysis: H. K., N. R.-Q., and C.-Y. L.

Funding acquisition: H. K. and C.-Y. L.

Investigation: H. K., N. R.-Q., X. H., A.J. E., L. A., and C.-Y. L.

Methodology: H. K., N. R.-Q., and C.-Y. L.

Project administration: H. K. and C.-Y. L.

Supervision: H. K. and C.-Y. L.

Visualization: H. K. and C.-Y. L.

Writing-original draft: H. K. and C.-Y. L.

Writing-review & editing: H. K. and C.-Y. L.

## Supplemental Figure Legends

**Supplemental Figure 1.**
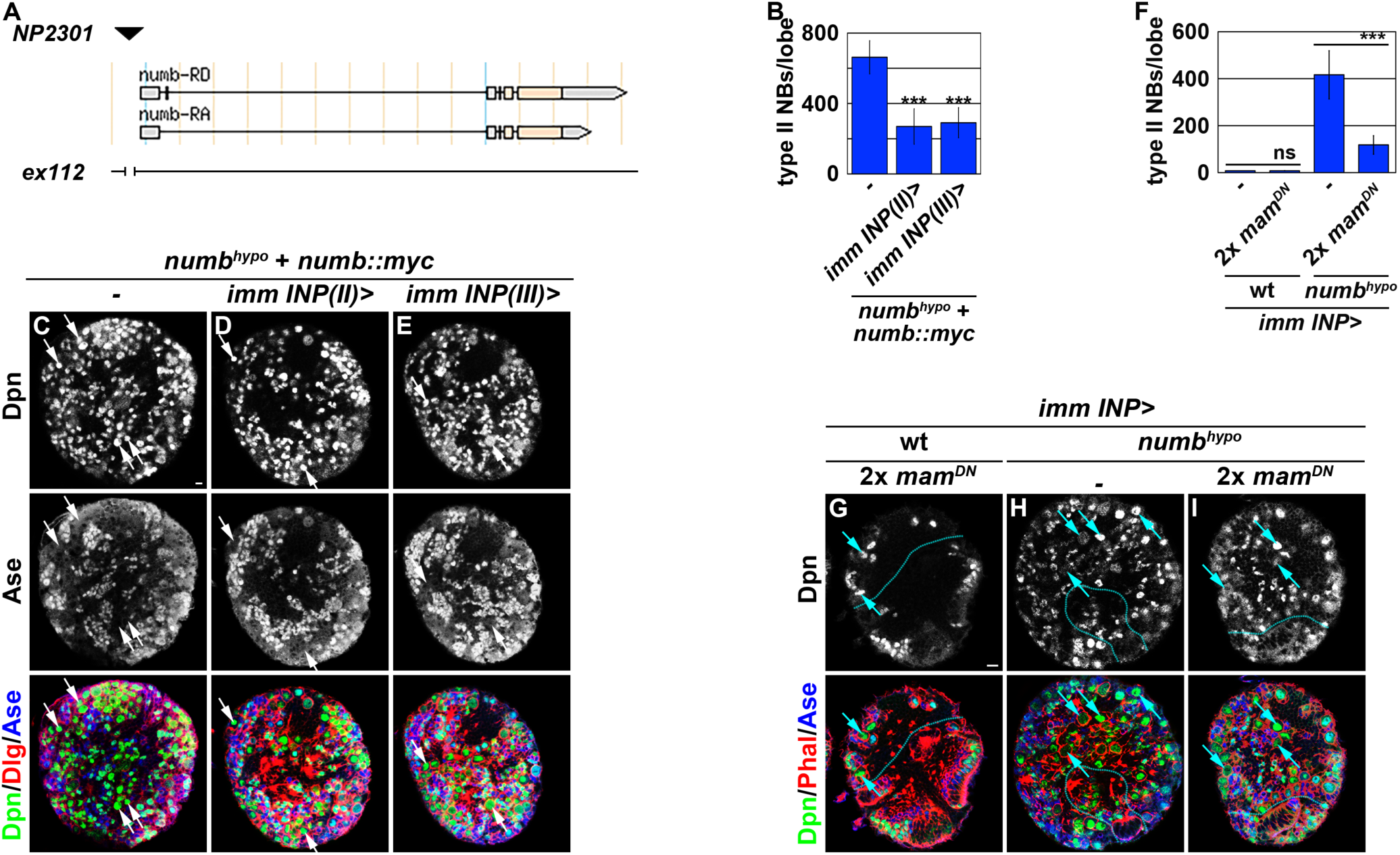
Generation and characterization of the *numb*^*hypo*^ allele. (A) The map shows the *numb*^*NP2301*^ allele induced by a transposable P-element insertion in the 5’-regulatory region of *numb*. (B-F) Restoring Numb::Myc expression in immature INPs rescued the supernumerary neuroblast phenotype in *numb*^*hypo*^ brains. The average number of type II neuroblasts per *numb*^*hypo*^ brain lobe with or without restoring Numb::Myc expression in immature INPs is shown in B. (F-I) Inhibiting transcriptional activation of Notch target genes in immature INPs by overexpressing Mam^DN^ suppressed the supernumerary neuroblast phenotype in *numb*^*NP2301*^ brains. The average number of type II neuroblasts per *numb*^*hypo*^ brain lobe with or without Mam^DN^ overexpression is shown in F. White arrowhead indicates Ase^-^ immature INP (Dpn^-^Ase^-^). Bar graphs are represented as mean ± standard deviation. The *cul3*^*06430*^ allele was used in these experiments. P-values: *** <0.005. Scale bar, 10 μm.

**Supplemental Figure 2.**
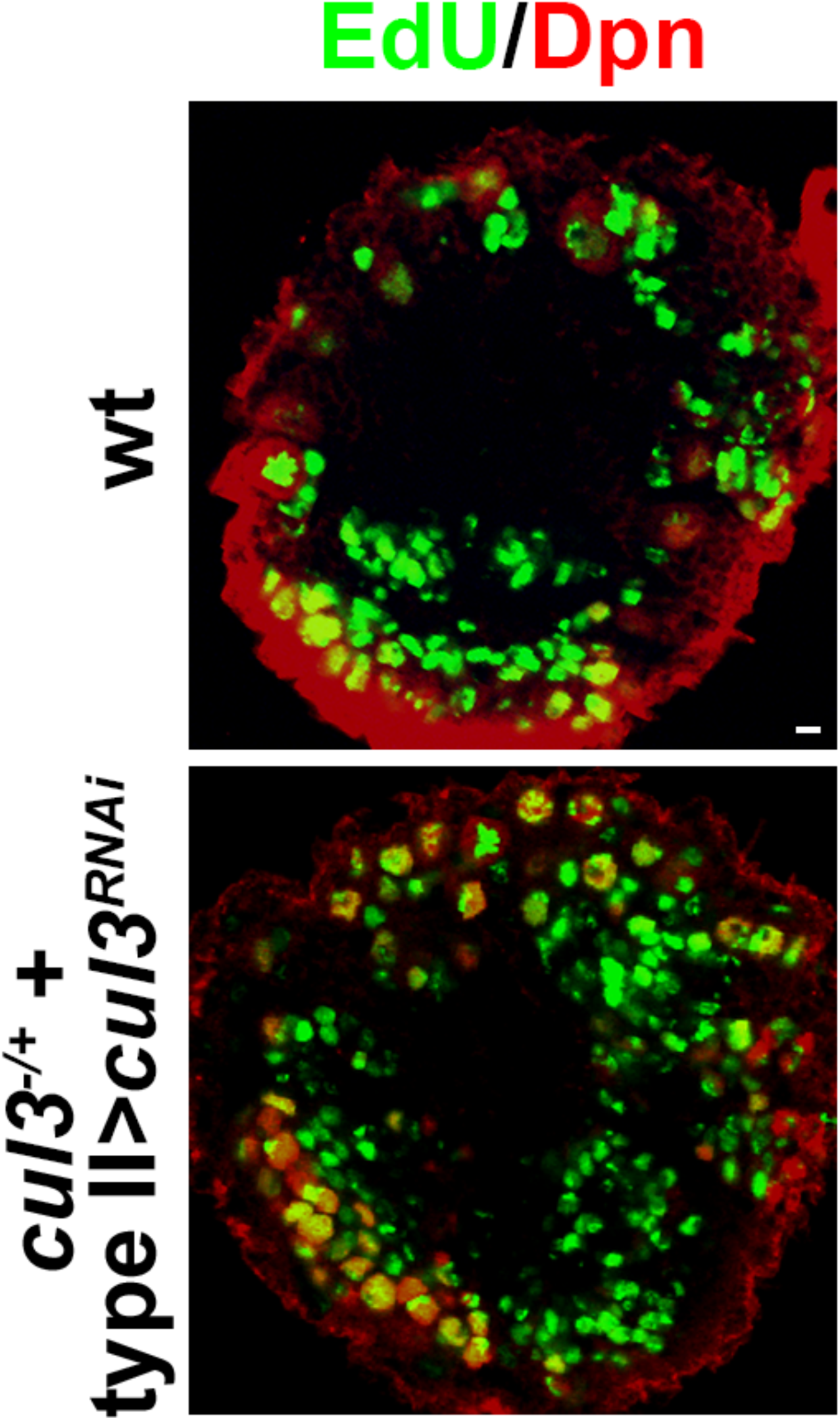
*cul3*-mutant brains display a supernumerary neuroblast phenotype. *cul3* heterozygous brains with endogenous *cul3* knocked down by RNAi showed a supernumerary neuroblast phenotype. Scale bar, 10 μm.

